# Mangrove forest structure and composition along urban gradients in Puerto Rico

**DOI:** 10.1101/504928

**Authors:** Benjamin Branoff, Sebastián Martinuzzi

## Abstract

Urban forests are repeatedly characterized as distinct in composition and structure in comparison with their non-urban counterparts. This holds true for mangroves, although previous studies lack quantified representations of urbanness as well as any inclusion of hydrology or water chemistry, which are important influences on mangrove forest structure, composition, and function. This study uses LiDAR and ground-based measurements of mangroves within well quantified urban gradients in Puerto Rico to test for the relative importance of urbanization alongside flooding metrics and surface water chemistry in explaining observed patterns of forest structure and composition. In simple regression, urban metrics were the most powerful predictors of forest composition but not structure. Results show higher tree diversity but lower mangrove diversity in the most urban forests. Structural measurements, however, were best explained by flooding, surface water chemistry, and non-urban land cover metrics. Nitrogen concentrations best explained stem density and tree size, while flooding metrics best explained stand biomass and basal area, and surrounding vegetation cover best explained canopy cover and height metrics. In multiple regression, land cover and surface water chemistry were more important than flooding, with population density again being the most important variable in explaining mangrove forest diversity. Results show that urbanization is an important influence on mangrove composition and basal area, leading to higher tree diversity and lower basal area, consistent with patterns in terrestrial forests. But urban mangrove forests are also lower in mangrove diversity and tend to have representation only by *Laguncularia racemosa*. Nitrogen concentrations and surrounding vegetation cover, both of which are indirectly influenced by urbanization, were positively related to tree size and canopy cover and height, respectively. These tests suggest urbanization is an important influence on mangrove forest structure and composition, but that flooding and water chemistry must also be considered when managing these forests.

## Introduction

Urbanization has been an important contributor to forest disturbance at the turn of the twentieth century, with trends indicating an increasingly important role in tropical coastal areas over the next few decades (DeFries, Rudel, Uriarte, & Hansen, 2010; Geist & Lambin, 2002; Jorgenson & Burns, 2007; Seto, Güneralp, & Hutyra, 2012). The conversion of forests to developed lands usually follows economic transitions that favor industrialized societies and continuing transitions may in some cases support regrowth, albeit with novel forests (Wright, 2005; Wright & Muller-Landau, 2006). As a result, forests in urban and surrounding lands are often represented by unique anthropogenic influences and ecological traits (Grimm et al., 2008; Pickett et al., 2008; Rowntree, 1984).

Characterizations of urban forests often consist of lower stem densities and larger individuals, with stands exhibiting increased edge openness and regeneration failure (Pickett et al., 2001). These forests are further generalized as having higher floral diversity, usually due to non-native species from residential gardens and municipal landscaping. As a result, the function of these systems is also novel as evident in altered community composition, biogeochemistry, productivity, and resiliency (Alberti, 2005; McDonnell & Pickett, 1990). This in turn, gives way to an adjusted provisioning and valuation of ecosystem services (Dwyer, McPherson, Schroeder, & Rowntree, 1992; McPherson et al., 1997). Much of this is known from the study of terrestrial systems, with comparatively little understood of the dominant forested biome of tropical and sub-tropical shorelines, mangroves.

Globally, mangrove coverage in the largest cities is decreasing faster than overall rates from corresponding countries (B. L. Branoff, 2017). In some cases, this loss results in fragmented forests consisting of novel species assemblages and size classes (Benfield, Guzman, & Mair, 2005; Mohamed, Neukermans, Kairo, Dahdouh-Guebas, & Koedam, 2009; Nortey, Aheto, Blay, Jonah, & Asare, 2016; Zamprogno et al., 2016). There are cases of expanding urban mangrove coverage (DasGupta & Shaw, 2013; Pham & Yoshino, 2016), although the young forests may contain less biomass (Friess, Richards, & Phang, 2015). These studies suggest a systemic influence of urban land-use on mangrove structure and composition, but most have been conducted with little or no quantification of urbanness, relying instead on qualitative definitions of urban and non-urban. Most were also done independently of hydrology or water chemistry, which have long been recognized as powerful influences on the same structural and compositional metrics (Lugo & Snedaker, 1974; Wolanski, Mazda, & Ridd, 1993).

Flooding metrics of hydroperiod and flood frequency are important for mangrove seedling survival and adult growth (Krauss, Doyle, Twilley, Rivera-Monroy, & Sullivan, 2006; McKee, 1995). Water salinity, pH, dissolved oxygen, phosphorus, and nitrogen concentrations are also known to influence mangrove physiology and forest structure and composition (Cardona & Botero, 1998; Feller, Whigham, McKee, & Lovelock, 2003; Joshi & Ghose, 2003; McKee, 1996; Reef, Feller, & Lovelock, 2010). Again, while there are some anecdotal connections between mangrove hydrology, water chemistry, and urbanization (Marois & Mitsch, 2017; Seguinot Barbosa, 1996; Tian-Hong, Peng, & Zhi-Jie, 2008), there remains no quantified connection between these components, leaving little definitive evidence for the hypothesis that urbanization influences mangrove forests through hydrology and water chemistry.

In Puerto Rico, Brandeis et al. (2014) provide a limited structural inventory of the mangrove and non-mangrove forests of San Juan based on a small sample area, but there are no conclusions regarding the influence of urban areas on this structure, and much less is known of the urban mangroves outside of the San Juan metropolitan area. Overall, urban mangrove coverage in Puerto Rico has largely failed to expand during the last quarter of the twentieth century, despite a 12% increase in mangroves across the island (Martinuzzi, Gould, Lugo, & Medina, 2009). Further, urban forests were found to have fewer and smaller forest fragments compared to more rural sites. Another study has shown variability in the flooding dynamics and water chemistry of the mangroves of Puerto Rico, but little influence from surrounding urbanization (Branoff, 2018). Thus, while there is some information on the composition of the urban mangroves of Puerto Rico, there is little evidence tying any observed changes in mangrove ecology to specific components of urban landscapes.

This study uses a combination of ground-based and remote sensing measurements to address the following objectives: 1) Characterize the structure and composition of mangrove forests along urban gradients in three watersheds of Puerto Rico, 2) Test for correlations between the measured forest characteristics and the three potential influences of urbanization, hydrolny, and water chemistry, and 3) Test for the combined importance of all three influences on mangrove forests through multiple regression. Results will be used to inform local management strategies as well as in future planned assessments of community ecolny and the provisioning of ecosystem services by these forests. Further, the Caribbean islands have been identified as one of five global biodiversity hotspots predicted to undergo a high urban area conversion before 2030 (Seto et al., 2012), presenting a particular challenge to the management and conservation of Caribbean mangroves (Ellison & Farnsworth, 1996; Lugo, Medina, & McGinley, 2014), and making this research especially important to generating informed management of both developed and non-developed lands in the region.

## Methods

### Study Location

Study sites were selected in Puerto Rico based on the range in urbanization surrounding their mangroves as described in (B. L. Branoff, 2018) (Figure 1). Urbanization for site selection was determined through the calculation of an urban index at each site. This index combines surrounding coverage of mangrove and non-mangrove vegetation, open water, and impervious surfaces, as well as population density and road length, into a single variable that is normalized from 0 to 100 to represent the least and most urban sites, respectively. Twenty one-hectare forested sites were established in three watersheds with the greatest range in urbanization: The San Juan Bay Estuary, the Río Inabón to the Río Loco (Ponce), and the Río la Plata (Levittown). All sites are bounded on one side by a water-body and most sites are roughly100 m by 100 m, with some asymmetry due to non-linear coastlines. In some cases (BAHMIN, BAHMAX, MPDMIN, MPNMIN, MPNMAX, TORMAX) forests were constricted by natural or anthropogenic features and these sites were extended along the shoreline to compensate for a lack of forest less than 100 meters from the shore. Human use of the mangroves is minimal at most sites, although harvesting of the crustacean, *Cardisoma guanhumi*, is sparsely evident at some and their use for grazing and corralling of horses, pigs, and dogs is also apparent. We found no evidence of wood harvesting at any of the sites.

**Figure 1.**
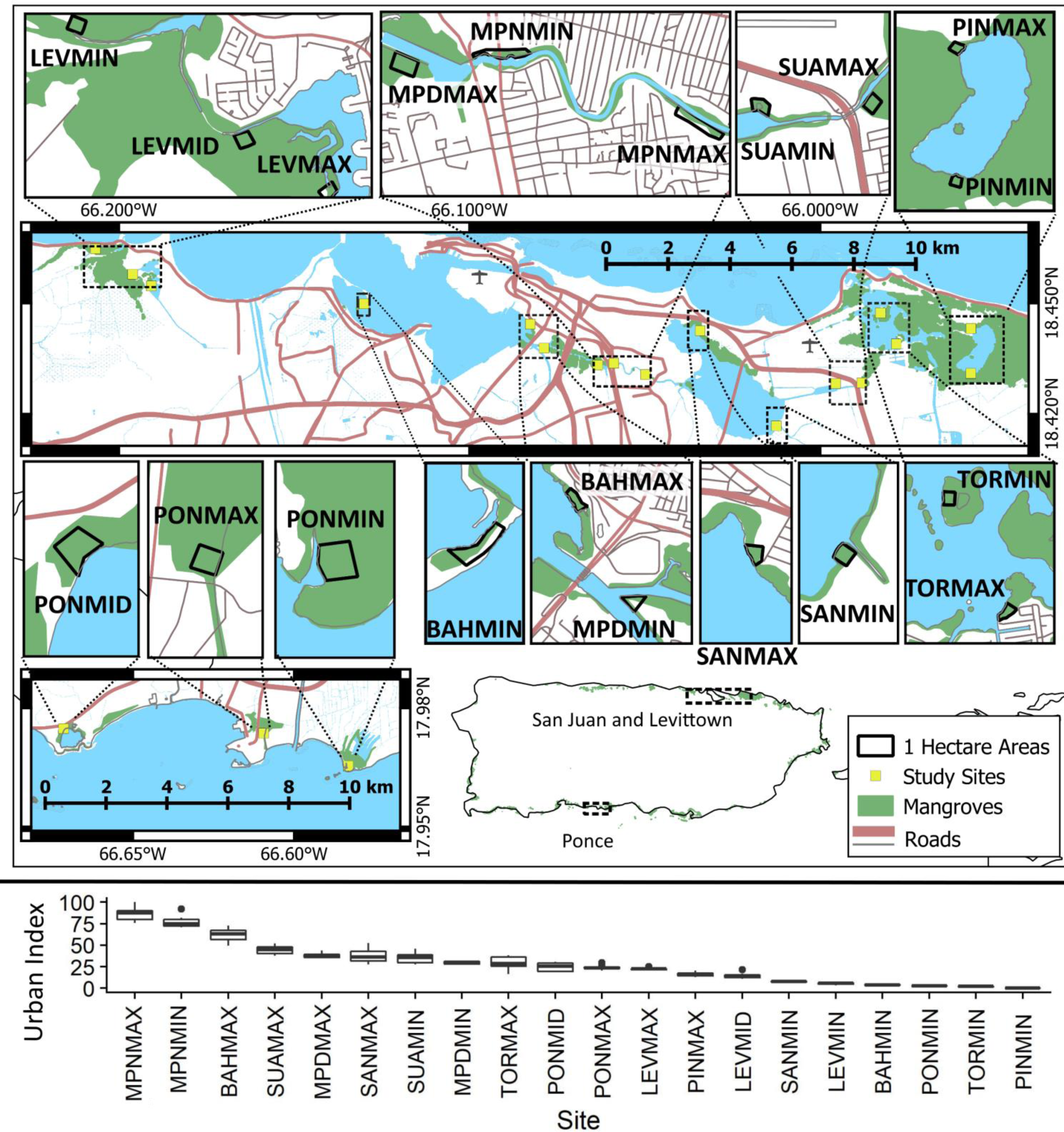
Study sites are twenty 1-hectare mangrove forests in three watersheds (top) and spanning a gradient of urbanization as measured by the urban index (bottom). The urban index is a combination of urban land use, vegetation and open water coverage, population density, and road length. A value of 0 represents the least urban site and a value of 100 is the most urban site. “MAX” and “MIN” postscripts refer to urbanization levels within each water body. BAH is the San Juan Bay, MPN is the Caño Martín Peña, SAN is the San José lagoon, SUA is the Suarez canal, TOR is the Torrecillas lagoon, PIN is the Piñones lagoon, LEV is Levittown and PON is Ponce

Fourteen sites were located throughout the roughly 1,500 ha of mangroves in The San Juan Bay Estuary. The mangroves were most recently described by Brandeis et al. (2014) as being dominated by *Rhizophora mangle*, although the authors point out this is likely due to sampling methods and that *Laguncularia racemosa* best characterize the forests and represents more biomass than any other species. Two sites were established in each of the seven primary waterbodies of the estuary, one each representing the minimum and maximum urbanized mangroves as described by (B. L. Branoff, 2018). Three sites each were established in the other two watersheds represented by the cities of Levittown and Ponce. Very little has been published on the mangroves of Levittown, which surround an estuary composed primarily of an artificial tidal lagoon constructed to drain surrounding settlements and connected to the ocean through a tidal creek and permanent inlet. Unlike the other two sites, the mangroves at Ponce are largely unconnected and do not share the same estuarine conditions. Additionally, Ponce receives a much drier southern climate in comparison to the northern sites, with a median annual rainfall of 755 mm versus 1600 mm on the northeastern coast (Branoff, 2018). Like Levittown, very little is known on the mangroves in Ponce although one study did characterize the hydrology of Punta Cabullones (Rodríguez-Martínez & Soler-López, 2014).

### Calculating Urbanization

Several spatial datasets were used to characterize the urbanization at each site and to extract variables for statistical testing (Table 1). All spatial analyses were performed in the R programming language (Yan *et al*., 2011) with the packages sp (R. S. Bivand, Pebesma, & Gomez-Rubio, 2013), rgeos (R. Bivand & Rundel, 2017), and raster (Hijmans, 2016). A repository of the R code used for the spatial sampling can be found at https://github.com/BBranoff. To begin, ten points were randomly placed in the one-hectare areas and served as the measurement replicates for each site. Point locations were determined randomly through the *spsample* function of the sp package. For each point, land cover and urbanization variables were extracted from sampling circles of radius 50 m, 100 m, 200 m, 500 m, and 1 km, as generated using the *gBuffer* function from rgeos.

**Table 1.** Spatial datasets used to determine relative elevations and to quantify the urbanization surrounding each study site. Variables were sampled from a sampling area described by a circle of radius 500 m and centered on individual study sites.

| Variable | Dataset Description | Source |
| --- | --- | --- |
| **Land Cover** |  |  |
| **Urban Vegetation and Water** | 2 m resolution land cover raster for Puerto Rico in 2010 | (Office for Coastal Management, 2017) |
| **Mangrove Coverage** | 30 m resolution continuous mangrove coverage raster for 2012 | (Hamilton & Casey, 2016) |
| **Population Density** | Total population shapefile for 2010 in Puerto Rico by census block | (U.S. Census Bureau, 2010) |
| **Road Length** | Road network shapefile for Puerto Rico in 2015 | (U.S. Census Bureau, 2015) |
|  | State highway system for Puerto Rico in 2015 | (Autoridad de carreteras y Transportación, 2015) |

Vegetation, Mangrove, Water, and Urban coverage were extracted from these circles by masking the corresponding rasters with the sampling circle through the *mask* function from the raster package, leaving only values within the circle. These values were then gathered through the *getValues* function also from the raster package. The mangrove raster contains values of continuous mangrove coverage within each cell, so total mangrove coverage was the sum of these values. The land cover raster, however, contains values indicating the dominant cover type within each cell. Thus, the coverage for each category of vegetation, water, and urban, was calculated by first summing the number of cells within each category and then dividing by the cell size. Urban coverage was associated with classes 2 and 5 (Impervious and Developed Open Space, respectively), vegetation was classes 6 through 18, and open water was class 21. The cell size was determined by taking the area of the circle over the total number of cells within the circle.

Road and highway lengths within the sampling circle were determined by first performing an intersect of the road network and the sampling circle through the *gIntersection* function from the rgeos package, leaving only the portions of roads within the circle. The lengths of all road segments were then summed through the *SpatialLinesLengths* function from the sp package. Population density was calculating using the total population from census blocks and assumed all people lived in non-road impervious surfaces. The area of non-road impervious surfaces was obtained in each census block by first calculating the total impervious surface area (class 2) within each block as described in the previous paragraph and subtracting the area from roads. This was then used to calculate the population density per non-road impervious area, which was applied to the non-road impervious areas extracted from each portion of the census block contained within the sampling area. These variables were combined into the urban index as described above and all were used separately in hypothesis testing.

### Ground-Based Measurements

A 5 m radius plot design was used as recommended for small stem forests (C. Sparks, Masters, & Payton, 2002). Plots were established by fixing the end of a 5 m rope at the randomly generated coordinates described above and extending the rope until tight. These center points were located using a Garmin eTrex 10 global positioning system with an accuracy of ± 3m. All woody plants greater than 1 cm in diameter at the breast and within the 5 m radius were identified and their diameter measured with a diameter measuring tape at a height of 1.4 m, or diameter at breast height (dbh). For individuals of *R*. *mangle* whose primary prop roots joined the trunk at heights greater than 1.4 m, trunk diameter was measured just above the confluence of the prop roots where a true mainstem existed. Branches from mainstems of all species were also measured if they originated at heights less than 1.4 m and if their dbh also exceeded 1 cm. These were categorized as side stems and the largest individual of a related group of stems was categorized as the mainstem.

Additional variables describing site hydrolny and water quality were also used in hypothesis testing and were derived from (B. L. Branoff, 2018). These include mean flooding dynamics metrics derived from five-year models of water levels at all sites. These metrics are the average depth (m), proportion of time flooded (fraction), mean daily flood frequency (d^-1^), mean flood length (days), number of days with at least one flood per year (days), flooded hours per year (hours) and dry hours per year (dry). Water chemistry measurements were made in surface waters within 1 km of each site on a monthly and bi-annual basis over the same five-year period by the San Juan Bay Estuary Program. These measurements were only available for sites in San Juan. Monthly measurements are dissolved oxygen (mg/L), pH, salinity (PSS), specific conductivity (ms/cm), and temperature (°C). Bi-annual measurements were Ammonia (mg/L), total Kjeldahl nitrogen (mg/L), nitrate & nitrite (mg/L), and phosphorus (mg/L).

### LiDAR

LiDAR was flown over the San Juan Bay Estuary sites in March of 2017 by the NASA G-LiHT program (Cook et al., 2013). Point clouds were processed at the one-hectare level, including all ten plots within a one-hectare forest. Tree height was measured in quantiles of 10% increments, from the 10^th^ to the 90^th^, representing the heights in which 10% and 90% of all trees are shorter, respectively. Canopy density was also measured for the same height deciles, and total canopy cover was the percentage of all first returns intercepted by a tree. The standard deviation of tree heights was also recorded, as was the skewness and kurtosis of the heights.

### Analyses and Statistical Testing

All numerical analyses and statistical tests were performed in R (Yan et al., 2011). Biomass estimations were calculated for each tree through allometric equations using the measured dbh and wood specific gravities when available. For the three true mangrove species, equations were species and dbh specific as derived from three separate sources on Caribbean mangroves (Fromard et al., 1998; Imbert, 1989; Smith & Whelan, 2006). The mean of these three values was used and when no value was available for greater size classes, a general equation for mangrove habitats was used from Chave et al. (2005). This equation was also used for non-mangrove species in combination with specific gravities derived from Reyes et al. (1992).

Statistical tests were performed at the site level using values from each plot, and at the watershed level by summing the various metrics from each plot at each site. Comparisons across sites and watersheds were done through an ANOVA as calculated by the *aov* function (Yan et al., 2011), and pairwise tests were performed through a Tukey Honest Significant Differences test as computed through the *TukeyHSD* function. Due to the high number of variables involved in the analysis, non-metric multidimensional scaling (NMDS) was first performed to observe overall patterns between forest metrics and urbanization. This was done twice, once on ground-based measurements and again on LiDAR measurements. Functions for this analysis are from the vegan package (Oksanen et al., 2018). NMDS was first performed through the *metaMDS* function on the ground-based and LiDAR derived variables. Results were then rotated through the *MDSrotate* function so that the horizontal axis of the NMDS was aligned with the vector describing the urban index. Vectors describing the ground-based and LiDAR measurements were then calculated through the *envfit* function, which finds the projection of the maximum correlation between each variable and the ordination. Finally, ellipses describing the 95% standard error confidence area for each site were drawn using the *ordiellipse* function.

Simple regression was then performed between all response variables of forest metrics and all predictor variables of land cover/urbanization, flooding, and water chemistry. Models were constructed through the *lm* function (Yan et al., 2011) in the form y∼x and y∼ln(x). The highest performing models were selected as those with the highest R^2^ value whose p-value was lower than 0.05.

Multiple regression was performed by including one variable from each of the three potential influences of land cover, hydrolny, and water chemistry such that:

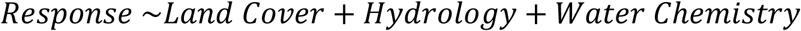

Models were constructed in the same way as simple regression, with both raw predictor variables as well as their natural logarithm. Bayesian Information Criteria (BIC) was calculated for all candidate models of the same response variable through the *glmulti* function of the same package name (Calcagno, 2013). BIC was used because it imposes a higher penalty than AIC for models with multiple predictors. The top three performing models for each response variable were selected based on the lowest BIC value and the significance of the model (p < 0.05). The relative importance of each variable to final models, as determined by its contribution to the overall R^2^, was calculated through the *calc*.*relimp* function of the relaimpo package (Grömping, 2006).

Graphs were produced through ggplot and the geom_violin, geom_bar, and geom_point functions for violin, bar and scatter plots, respectively (Wickham, 2009). Violin plots represent data at the plot level. Scatter plots represent data at the site level, and linear models were plotted through the stat_smooth function using the formula y∼x, unless it was outperformed by the formula y∼ln(x) as determined by the highest R^2^ value.

## Results

9,381 stems belonging to 7,250 mainstems were identified as belonging to a total of 30 different species, resulting in an overall stem density of 6,032 per hectare (Table 1). Fifty-one percent of these stems were represented by *Laguncularia racemosa*, 29% were *Rhizophora mangle*, 9% were *Avicennia germinans*, 7.5% were *Thespesia populnea*, 1% was *Calophyllum spp*., and the remaining 2.5% were made up of 25 other species. Basal area was 37.6 m^2^/ha on average, 63% of which was represented by *L*. *racemosa*, 21% by *R*. *mangle*, 7% by *A*. *germinans* and 9% by other species. The forests held 196 Mg/ha of woody biomass on average, with similar percentages by species as those of basal area. There were no significant differences in dbh, stem density, basal area, or biomass among the three watersheds (ANOVA; p = 0.5, 0.1 & 0.2, respectively).

At the site level, LEVMIN contained trees with a mean DBH on average 5.8 cm greater than trees at all other sites except MPDMAX, MPNMIN, MPNMAX, SANMIN, SANMAX, and TORMIN (ANOVA; p < 0.05) (Figure 2). LEVMID harbored 8,911 more stems per hectare on average than all other sites except BAHMIN and SUAMIN (ANOVA; p < 0.05). However, in basal area and biomass, only MPNMAX differed significantly and only from BAHMAX, PINMAX, PINMIN, and PONMIN, for which it held 30.8 m^2^/ha more basal area and 181 Mg/ha more biomass on average (ANOVA; p < 0.05). MPNMAX contained an average of three more species than all other sites except MPDMIN, which in turn held on average of two more species than LEVMIN, LEVMID, PONMAX and PONMIN (ANOVA; p < 0.05). Two urban sites were distinctly less diverse in mangrove diversity while two less-urban sites distinctly more diverse in mangrove species. LEVMAX and MPNMAX, both highly urban sites, contained the least number of species (one) and were statistically less diverse than half of the other sites (ANOVA; p < 0.05). PINMIN and PINMAX, two less-urban sites, were the most distinct in harboring high mangrove diversity at each plot, with an average of 2.6 and 2.5 species, respectively, and were significantly more diverse than seven of the other sites (ANOVA; p < 0.05).

**Figure 2.**
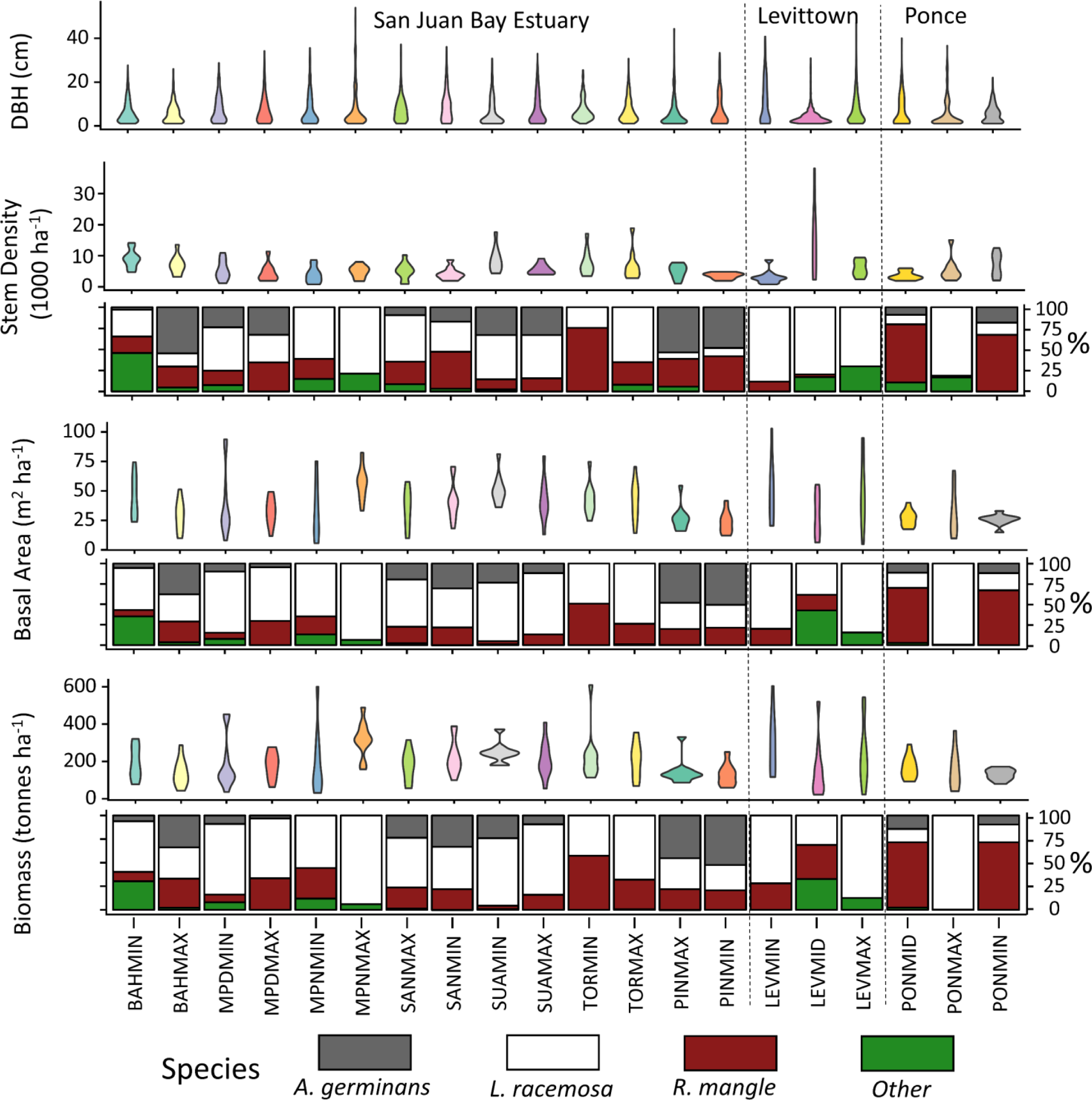
Forest structure and composition based on ground measurements. Bar plots show the relative proportion of the above violin plot from each group of species. Most sites are dominated by *L*. *racemosa*, which was the only mangrove species in some of the most urban sites.

LiDAR metrics were only available for sites in the San Juan Bay Estuary (Figure 3). The canopy height at BAHMIN was 6 meters lower on average than the other sites except BAHMAX, MPDMIN, SANMAX, TORMAX and TORMIN (ANOVA; p < 0.05). BAHMAX was 5 meters lower on average than MPDMAX, SANMIN, SUAMAX, PINMAX and PINMIN, and PINMIN was 6 meters higher on average than all sites except MPDMAX, MPNMAX, SANMIN, SUAMAX and PINMAX (ANOVA; p < 0.05). The standard deviation of all return heights was on average 2.3 meters greater at PINMIN than all other sites except PINMAX (ANOVA; p < 0.001). Mean canopy cover was 97% across all sites and did not vary among sites.

**Figure 3.**
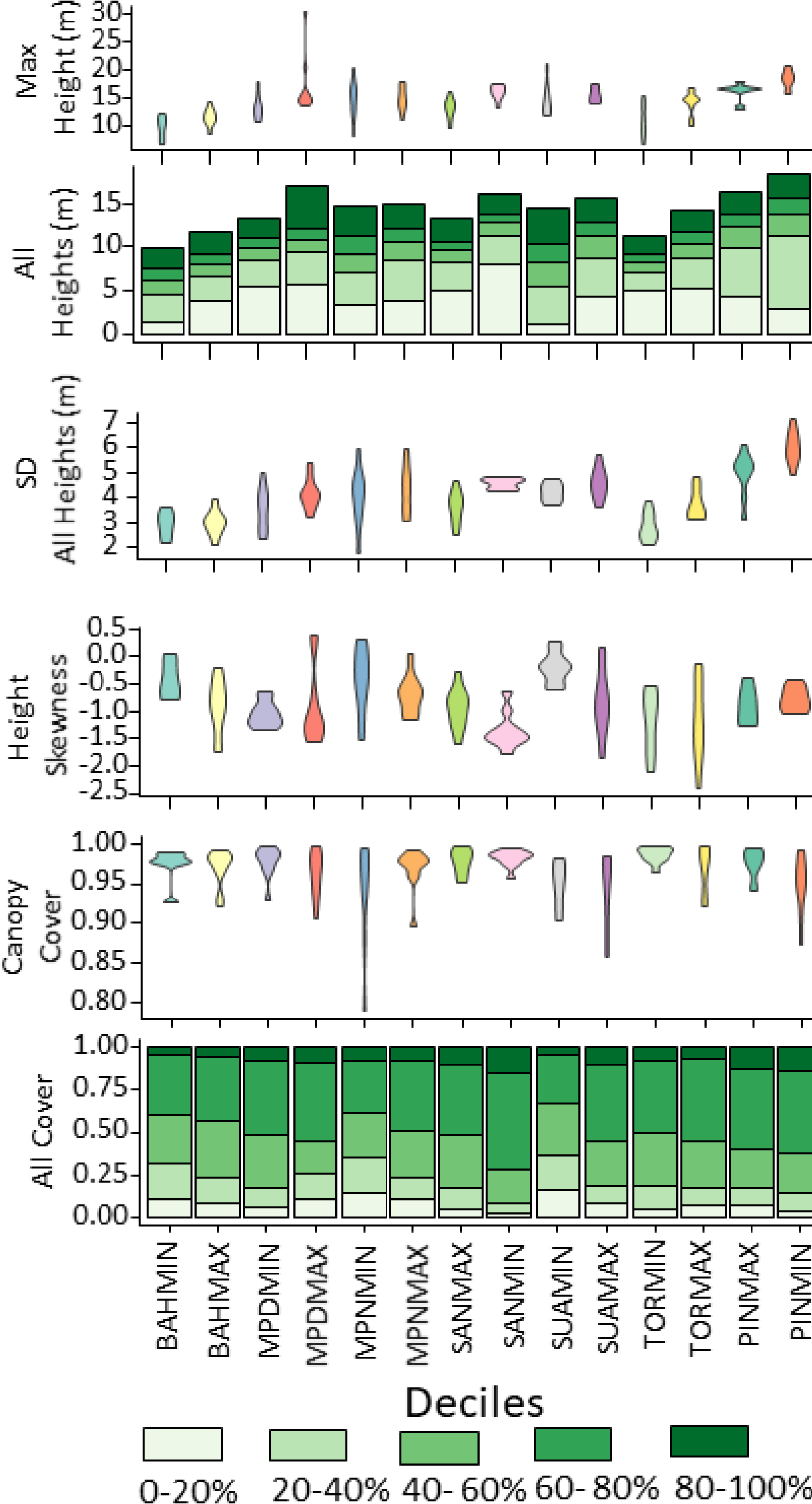
Forest structure based on LiDAR measurements. Bar plots of heights show the specific heights and covers of each decile group.

### Non-metric Multidimensional Scaling

Stress values in NMDS scaling were 0.14 for ground based measurements and 0.05 for LiDAR measurements, suggesting fair and excellent representation of actual data, respectively (Clarke & Ainsworth, 1993) (Figure 4). Further, all variables except tree diversity, were significantly correlated with the two NMDS axes (p < 0.001), suggesting their vectors are well represented in the ordination. Tree diversity was thus omitted from the ordination. Few sites were distinct from the others when represented by the ordination, however PINMIN is notable as having greater diversity, greater representation of *A*. *germinans*, and greater maximum height, as well as lower maximum DBH, basal area, and biomass. With the urban index aligned along the same axis as NMDS 1, variables whose absolute value of cosine(NMDS1) is greater than 0.5, are also aligned with the urban index. These are all variables except the percentage of biomass as *R*. *mangle*, the stem density, and the maximum tree height. Thus, most of the variables have some correlation along the urban gradient, although the strength and significance of these correlations cannot be inferred from the ordination. These correlations, among others, are tested in the following section.

**Figure 4.**
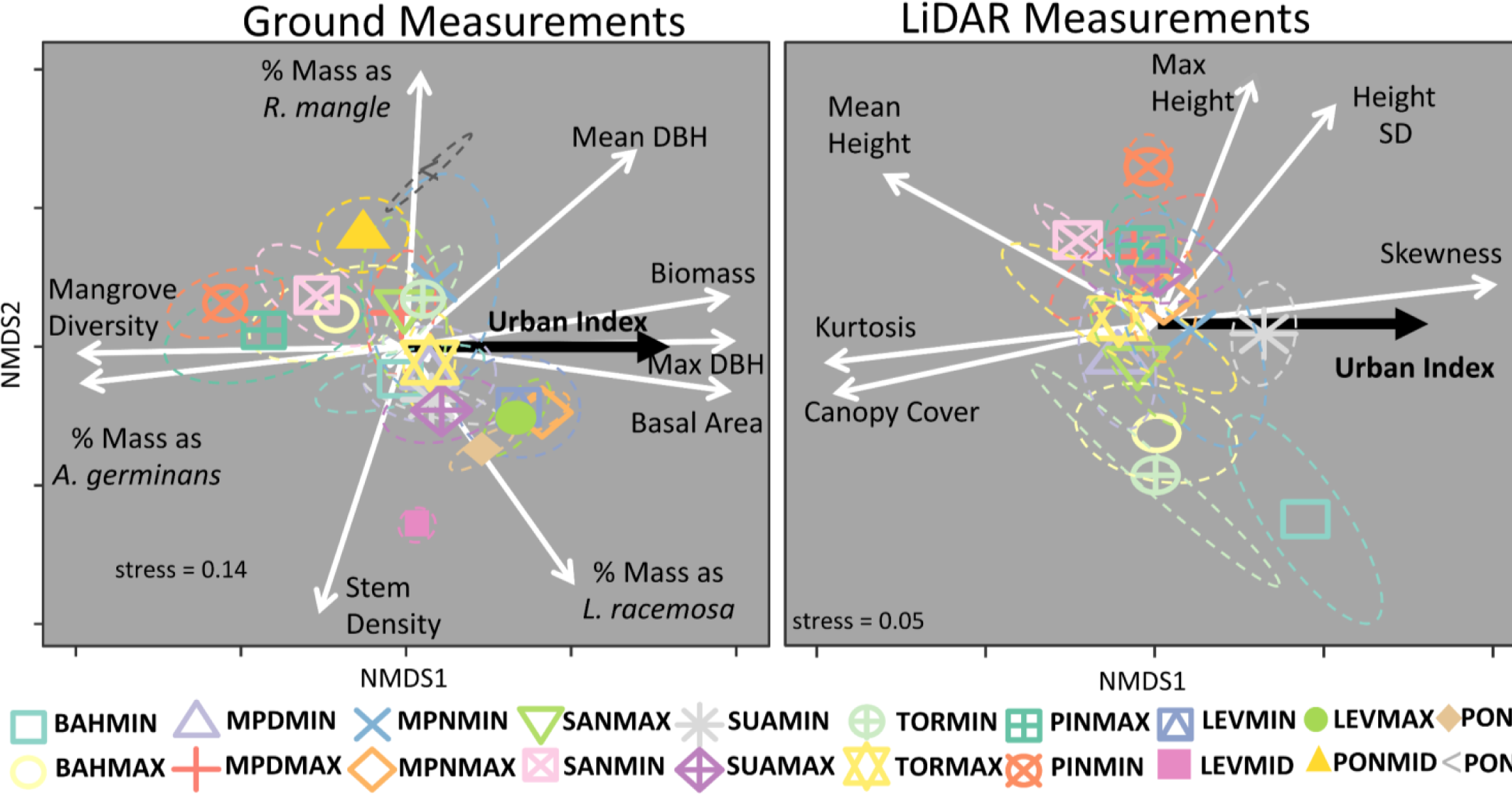
Non-metric multidimensional scaling of sites based on ground based (left) and LiDAR (right) measurements. Symbols are centroids of all plots at corresponding sites and ellipses are 95% standard error confidence ellipses for the location of centroids. Gray arrows are the directions of positive change for each structural and compositional measurement, and the black arrow is the direction of increasing urbanization based on the urban index. Only arrows for significant variables are shown. There is little separation of sites based on forest structure and composition, but some of the least urban sites (PINMIN, TORMIN, PONMIN, and BAHMIN) are most unique in comparison to the other sites as indicated by the separation of their ellipses from the others. Roughly half of the variables are partially aligned along the urban index axis, indicating a potential correlation with urbanness.

### Linear models

In linear regression, the strongest models involving urban variables were those describing forest composition, while forest structural metrics involved primarily flooding and water chemistry predictors (Figure 5). The percent of mangrove biomass as *A*. *germinans* decreased with the surrounding urban index within 1 km (R^2^ = 0.52, p < 0.001), and the percent of mangrove biomass as *L*. *racemosa* increased with the surrounding urban coverage also within 1 km (R^2^ = 0.34, p < 0.01). Also, the overall tree species diversity increased with surrounding population density within 200 m (R^2^ = 0.48, p < 0.001). The significance of this model, however, was due entirely to the relatively high species numbers at MPNMAX and could not be repeated when this site was removed from the test (R^2^ = 0.004, p = 0.3).

**Figure 5.**
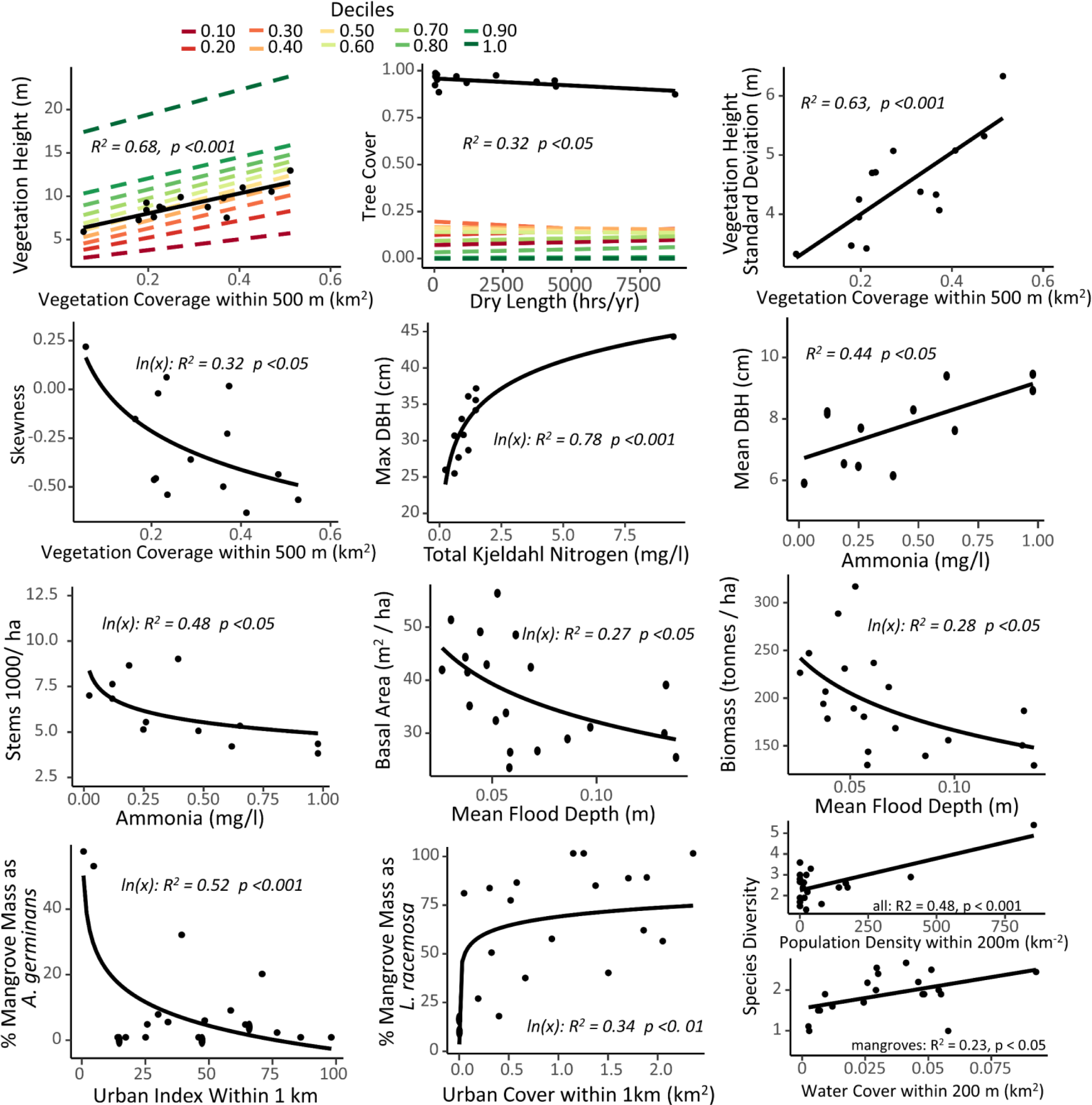
Forest structure and composition measurements with their strongest predictor variables from the analysis. Flooding dynamics, surface water nitrogen concentrations, and surrounding land cover are all strong predictors of individual forest structure and composition metrics.

The other response variables, mostly structural, were most strongly modeled using other metrics of land cover, hydrology, or water chemistry (Figure 5). Mean forest height, the standard deviation of the height, and the height skewness were all modeled with surrounding vegetation coverage within 500 m (R^2^ = 0.68, p < 0.001; R^2^ = 0.63, p < 0.001, R^2^ = 0.32, p < 0.05 respectively). Nitrogen was also a common strong predictor for the models. The natural logarithm of total Kjeldahl nitrogen was the strongest predictor for the maximum DBH at a site (R^2^ = 0.73, p < 0.001). Similarly, the natural logarithm of ammonia was the best predictor for stem density (R^2^ = 0.48, p < 0.05), and its raw form was best for predicting mean DBH (R^2^ = 0.44, p < 0.05). Flooding metrics were also common predictors. The natural logarithm of mean flood depth was the best predictor for stand basal area (R^2^ = 0.27, p < 0.05) and biomass (R^2^ = 0.28, p < 0.05), and the total dry duration was the best predictor for overall tree cover (R^2^ = 0.32, p < 0.05). Finally, the extent of surrounding water coverage was the best predictor of the number of mangrove species (R^2^ = 0.23, p < 0.05).

To test for the influence of multiple variables on forest structure and composition, candidate models for each response variable were constructed from combinations of one, two, or three variables from the groups of land cover, hydrology, and water chemistry metrics. Top ranked models by BIC almost always included all three variables and resulting R^2^ values averaged 0.68 (Table 2). In these models, land cover and water quality variables were the most important variables, averaging 46% and 42%, respectively, of the explanation of the variance (importance) in forest structure and composition. The importance of land cover and water quality was around 25% higher than flooding when predicting structural metrics (ANOVA; p < 0.001), but there was no difference for compositional metrics. Salinity had the highest mean importance, 72%, but appeared in only two models. Population density followed at 59% mean importance and appeared in seven models, and water coverage appeared in fourteen models and averaged 51% importance. Flood metric importance averaged about half of land cover and water chemistry, at 22% importance. The number of flood days was the most important flood variable, with an average importance of 39% and appearing in ten models. Nitrogen variables were the most common water chemistry variables, collectively appearing in thirty-seven models. Nitrate and Nitrite was the most important water chemistry variable on average, with a mean importance of 47%, and appeared in thirteen models. Among the land cover variables, fifty meters was the most common sampling distance, appearing in twenty of the models.

**Table 2.** Average and standard deviation of stem, basal area, and biomass per hectare of all species within the three watersheds included in the study.

|  | San Juan Bay Estuary |  |  | Levittown |  |  | Ponce |  |  | All |  |  |
| --- | --- | --- | --- | --- | --- | --- | --- | --- | --- | --- | --- | --- |
| Species | Stems / ha | Basal Area<br>m<sup>2</sup> / ha | Biomass<br>Mg / ha | Stems / ha | Basal Area<br>m<sup>2</sup> / ha | Biomass<br>Mg / ha | Stems / ha | Basal Area<br>m<sup>2</sup> / ha | Biomass<br>Mg / ha | Stems / ha | Basal Area<br>m<sup>2</sup> / ha | Biomass<br>Mg / ha |
| All | 5853 ± 1836.5 | 38.9 ± 9.5 | 200.2 ± 50.4 | 7724.3 ± 5724.4 | 42.6 ± 10.9 | 225.5 ± 69.7 | 5177.8 ± 1690.7 | 26.7 ± 1.3 | 146.3 ± 20.1 | 6032.4 ± 2578.6 | 37.6 ± 9.9 | 195.9 ± 53.2 |
| *Laguncularia racemosa* | 2572.8 ± 1704.8 | 24.3 ± 12 | 119.5 ± 62.2 | 6472.3 ± 6149.9 | 32.8 ± 10.1 | 165.5 ± 70.6 | 1884.4 ± 2470.5 | 11.9 ± 14 | 58.6 ± 71 | 3054.4 ± 2973.9 | 23.7 ± 12.9 | 117.3 ± 68 |
| *Rhizophora mangle* | 1893.3 ± 1546.1 | 8.4 ± 6.2 | 50.3 ± 38.1 | 169.8 ± 147 | 6.2 ± 1.1 | 55.8 ± 5.4 | 2915.7 ± 2982.3 | 13.8 ± 12 | 80.8 ± 70.4 | 1788.1 ± 1787.9 | 8.1 ± 7 | 50.4 ± 41.7 |
| *Avicennia germinans* | 709.4 ± 825.5 | 5.2 ± 5.1 | 27.6 ± 29.6 | 0 ± 0 | -- | -- | 220.7 ± 244.7 | 1.4 ± 0.4 | 9 ± 6.2 | 529.7 ± 745.4 | 2.7 ± 4.9 | 14.7 ± 27.8 |
| *Thespesia populnea* | 427.4 ± 1031.5 | 2.6 ± 5.1 | 9.8 ± 19.1 | 912.5 ± 1334.9 | 6.8 ± 6.2 | 26.5 ± 24.3 | 72.2 ± 125 | 0 ± -- | 0.1 ± -- | 446.9 ± 986.9 | 1.7 ± 5.1 | 6.6 ± 19.4 |
| *Calophyllum sp.* | 88.2 ± 288.1 | 0.7 ± 1.1 | 2 ± 3.3 | 38.2 ± 66.2 | 0 ± -- | 0.1 ± -- | -- | -- | -- | 67.5 ± 241.7 | 0.1 ± 1 | 0.3 ± 2.8 |
| *Dalbergia ecastaphyllum* | 26.4 ± 75.8 | 0.1 ± 0.1 | 0.3 ± 0.4 | 97.6 ± 169.1 | 0.4 ± -- | 1.6 ± -- | -- | -- | -- | 33.1 ± 88.3 | 0 ± 0.2 | 0.1 ± 0.7 |
| *Bucida buceras* | 32.7 ± 93.5 | 0.2 ± 0.2 | 1 ± 0.8 | -- | -- | -- | -- | -- | -- | 22.9 ± 78.9 | 0 ± 0.2 | 0.1 ± 0.8 |
| *unknown fabaceae* | 18.2 ± 64.5 | 0.9 ± 1.3 | 4.6 ± 6.4 | -- | -- | -- | -- | -- | -- | 12.7 ± 54 | 0.1 ± 1.3 | 0.5 ± 6.4 |
| *Jacquinia arborea* | -- | -- | -- | -- | -- | -- | 80.6 ± 139.7 | 0.1 ± -- | 0.3 ± -- | 12.1 ± 54.1 | 0 ± -- | 0 ± -- |
| *Roystonea borinquena* | 14.6 ± 31.1 | 1.7 ± 3.1 | 9.9 ± 19.9 | 4.2 ± 7.4 | 0.3 ± -- | 1.3 ± -- | -- | -- | -- | 10.8 ± 26.5 | 0.4 ± 2.9 | 2.5 ± 18.1 |
| *Schefflera morototoni* | 10 ± 33.9 | 0.2 ± 0.2 | 0.4 ± 0.5 | 4.2 ± 7.4 | 0 ± -- | 0 ± -- | -- | -- | -- | 7.6 ± 28.4 | 0 ± 0.2 | 0 ± 0.4 |
| *Terminalia catappa* | 10 ± 26.5 | 1.6 ± 0.4 | 8.4 ± 1.5 | 4.2 ± 7.4 | 0.3 ± -- | 1.2 ± -- | -- | -- | -- | 7.6 ± 22.4 | 0.2 ± 0.8 | 0.9 ± 4.3 |
| *Ardisia elliptica* | 10 ± 30.9 | 0 ± 0 | 0 ± 0 | -- | -- | -- | -- | -- | -- | 7 ± 26 | 0 ± 0 | 0 ± 0 |
| *Cocos nucifera* | 7.3 ± 17.8 | 1.5 ± 1.5 | 8.4 ± 8.9 | 8.5 ± 14.7 | 1.5 ± -- | 8.2 ± -- | -- | -- | -- | 6.4 ± 15.7 | 0.3 ± 1.2 | 1.7 ± 7.3 |
| *Mammea americana* | 4.5 ± 17 | 0 ± -- | 0.1 ± -- | -- | -- | -- | -- | -- | -- | 3.2 ± 14.2 | 0 ± -- | 0 ± -- |
| *Andira inermis* | 3.6 ± 13.6 | 0 ± -- | 0.1 ± -- | -- | -- | -- | -- | -- | -- | 2.5 ± 11.4 | 0 ± -- | 0 ± -- |
| *Coccoloba uvifera* | 3.6 ± 13.6 | 0 ± -- | 0.1 ± -- | -- | -- | -- | -- | -- | -- | 2.5 ± 11.4 | 0 ± -- | 0 ± -- |
| *Tabebuia heterophylla* | 3.6 ± 13.6 | 0 ± -- | 0 ± -- | -- | -- | -- | -- | -- | -- | 2.5 ± 11.4 | 0 ± -- | 0 ± -- |
| *Annona glabra* | -- | -- | -- | 12.7 ± 22.1 | 0.5 ± -- | 2.7 ± -- | -- | -- | -- | 1.9 ± 8.5 | 0 ± -- | 0.1 ± -- |
| *Conocarpus erectus* | 1.8 ± 6.8 | 1.1 ± -- | 7.6 ± -- | -- | -- | -- | 4.2 ± 7.4 | 0.4 ± -- | 2.3 ± -- | 1.9 ± 6.2 | 0.1 ± 0.5 | 0.5 ± 3.7 |
| *Hippomane mancinella* | 2.7 ± 10.2 | 0 ± -- | 0 ± -- | -- | -- | -- | -- | -- | -- | 1.9 ± 8.5 | 0 ± -- | 0 ± -- |
| *Pavonia fruticosa* | 2.7 ± 10.2 | 0 ± -- | 0 ± -- | -- | -- | -- | -- | -- | -- | 1.9 ± 8.5 | 0 ± -- | 0 ± -- |
| *Puallinia pinnata* | 1.8 ± 4.6 | 0.1 ± 0.1 | 0.5 ± 0.6 | -- | -- | -- | -- | -- | -- | 1.3 ± 3.9 | 0 ± 0.1 | 0 ± 0.6 |
| *Unknown 1* | 1.8 ± 6.8 | 0.1 ± -- | 0.3 ± -- | -- | -- | -- | -- | -- | -- | 1.3 ± 5.7 | 0 ± -- | 0 ± -- |
| *unknown palm* | 1.8 ± 6.8 | 0 ± -- | 0.1 ± -- | -- | -- | -- | -- | -- | -- | 1.3 ± 5.7 | 0 ± -- | 0 ± -- |
| *Unknown 2* | 0.9 ± 3.4 | 0 ± -- | 0 ± -- | -- | -- | -- | -- | -- | -- | 0.6 ± 2.8 | 0 ± -- | 0 ± -- |
| *Unknown3* | 0.9 ± 3.4 | 0 ± -- | 0 ± -- | -- | -- | -- | -- | -- | -- | 0.6 ± 2.8 | 0 ± -- | 0 ± -- |
| *Unknown 4* | 0.9 ± 3.4 | 0 ± -- | 0 ± -- | -- | -- | -- | -- | -- | -- | 0.6 ± 2.8 | 0 ± -- | 0 ± -- |
| *Unknown 5* | 0.9 ± 3.4 | 0 ± -- | 0 ± -- | -- | -- | -- | -- | -- | -- | 0.6 ± 2.8 | 0 ± -- | 0 ± -- |
| *Unknown 6* | 0.9 ± 3.4 | 0 ± -- | 0 ± -- | -- | -- | -- | -- | -- | -- | 0.6 ± 2.8 | 0 ± -- | 0 ± -- |

## Discussion

Results from three separate statistical methods confirm a significant influence of urbanization on mangrove forest composition and structure, but that this influence is shared between flooding dynamics and water chemistry. Overall, land cover and water chemistry metrics were 20% more important than flooding metrics in explaining the variation in forest structure. Linear models with individual variables showed that urban variables were strongest in predicting compositional responses of forest metrics, and multiple regression models confirmed this. This suggests that urbanization is indeed influencing forest composition and structure, but hydrology and water chemistry should also be considered for a more complete understanding of urban mangrove forests.

Tree girths, as indicated by the maximum and mean diameters at each site, were most strongly predicted by surface water nitrogen in both simple (Figure 5) and multiple regression (Table 2). Much of the remaining variation in the data was explained by the surrounding road length or population density (Table 2). This agrees with urban areas in general, which typically contain a greater percentage of large trees than do less urban forests (Nowak & Crane, 2002). The reason for this varies and may be due to municipal bans on large tree removal (Mynors, 2002) as well as increased temperature and CO2 concentrations in urban areas (Pretzsch et al., 2017). Mangroves have been observed to grow larger under enriched CO2 conditions (Farnsworth, Ellison, & Gong, 1996). But urban mangroves are also distinct from terrestrial urban forests in that they are permanently or periodically flooded by urban waters, which likely carry elevated nitrogen concentrations that have been tied to roads and sewage (Bettez, Marino, Howarth, & Davidson, 2013; Brush, 2009; Davidson, Savage, Bettez, Marino, & Howarth, 2010). Such enrichment of nitrogen has been shown to increase girth of mangrove seedlings (Agraz-Hernández, del Río-Rodríguez, Chan-Keb, Osti-Saenz, & Muñiz-Salazar, 2018), although no studies were found linking maximum diameter to nitrogen. Thus, given the multiple regression results, its likely urban sourced nitrogen, especially from roads and sewage, is driving an increase in maximum tree girths in the more urban forests.

**Table 2.**
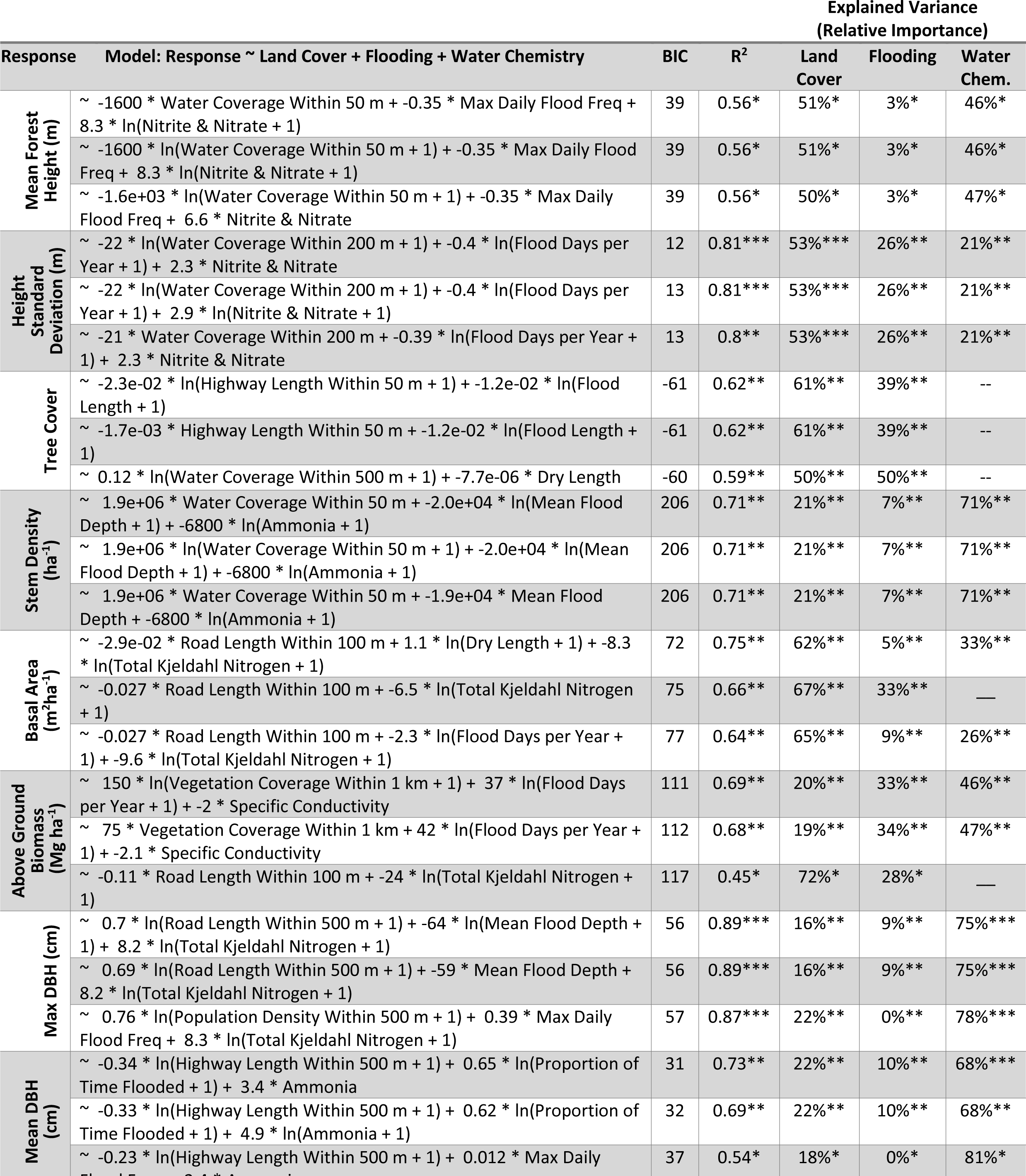
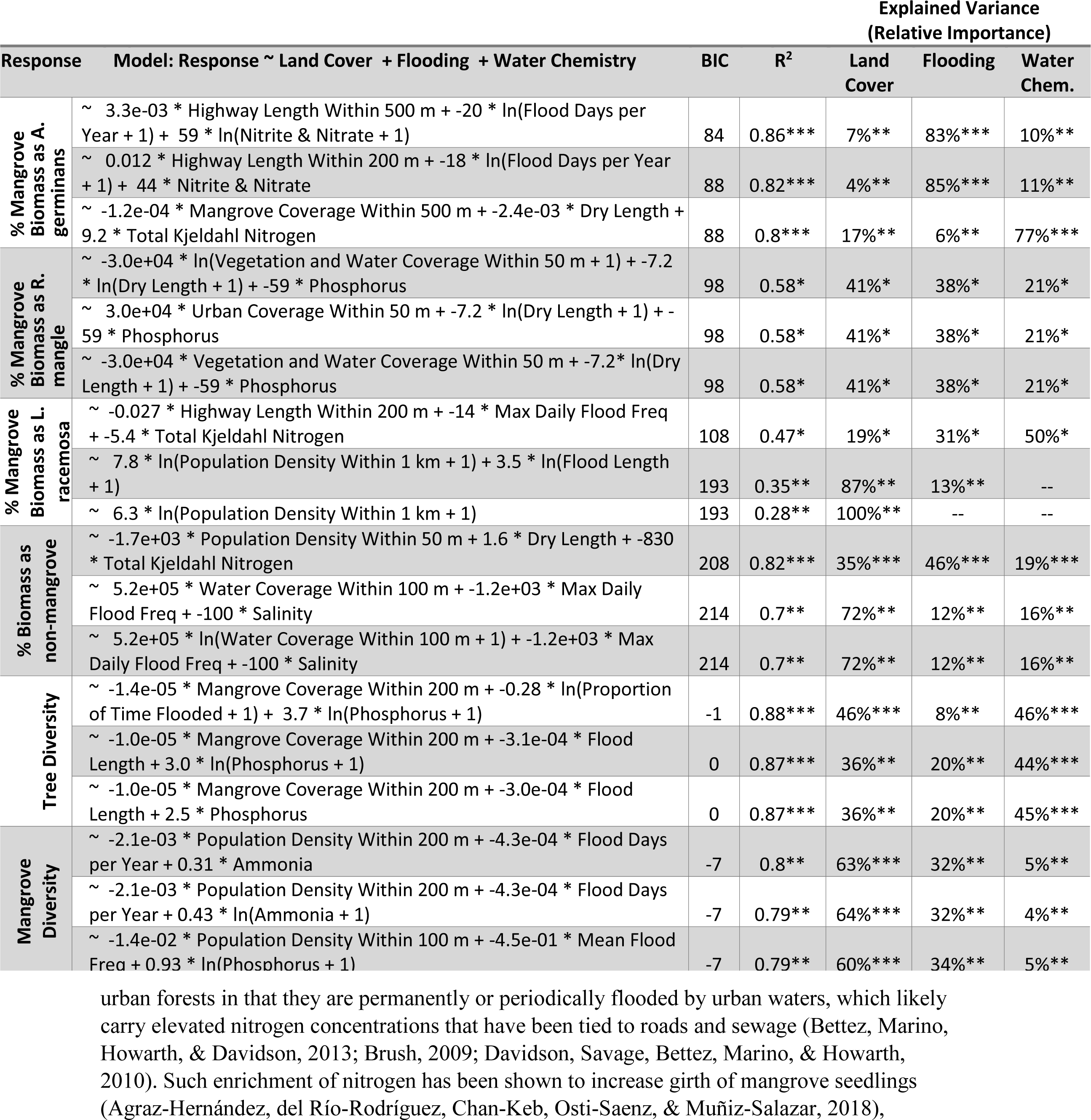
Top models for forest structure metrics as determined by the lowest BIC for models containing all combinations of surrounding land cover, hydrology, and water quality. Asterisks are p values of * <0.05, **<0.01 & ***<0.001. Top models for forest composition metrics as determined by the lowest BIC for models containing all combinations of surrounding land cover, hydrology, and water quality.

Tree height was also strongly explained by nitrogen concentrations but not by any explicit urban variable. The most explanatory variable in describing vegetation height metrics in both simple and multiple regression, was the surrounding extent of either open water or vegetation cover, which could also be taken as the absence of any urban area. Still, there were no significant correlations between urban variables and the height metrics. This agrees with an overall lack of any study describing such a relationship in other urban forests, although there are studies that attribute mangrove height to soil organic matter (Costa, Dórea, Mariano-Neto, & Barros, 2015), salinity (Lin & Sternberg, 1992), or combinations of nutrient availability and stress (Feller et al., 2003; McKee, Feller, Popp, & Wanek, 2002). We found no relationship between forest height and surface water salinity, and we did not measure pore-water conditions, but the observed relationship with surface water nitrogen concentrations agrees with previous studies. The influence of surrounding water and vegetation coverage on our height models may be explained by the persistent pressure from tropical storms, which is known to limit the height of Caribbean mangroves (Pool, Snedaker, & Lugo, 1977). Our models suggest a negative relationship between height and surrounding open water, and a positive relationship between height and surrounding vegetation. This may imply that higher wind speeds and storm surge associated with open water forests may be imposing a greater height pressure on trees in comparison to those in more inland forests.

Overall forest structure, as measured by stem density, basal area, and biomass, was most strongly correlated with nitrogen and flooding metrics in simple regression (Figure 5), but importance was mixed in multiple regression (Table 2). Urbanization was strongest in the basal area model, where road density had a negative effect. Similarly, stem density and biomass were positively related to water and vegetation coverage, which again could be taken as the absence of urban area. A number of studies have reported lower stem densities and basal areas in urban mangrove forests (Mohamed et al., 2009; Nagendra, Sudhira, Katti, & Schewenius, 2013; Nfotabong-Atheull, Din, & Dahdouh-Guebas, 2013), although these are primarily associated with wood harvesting, which is not a significantly common practice in Puerto Rico. Another study has reported lower urban mangrove biomass independent of harvesting and attributes this to the age and successional state of the forest (Friess et al., 2015). This could be the case for the urban mangroves of San Juan as well, which experienced a period of deforestation during the 1960s, and are perhaps still recovering (Martinuzzi et al., 2009). Thus, despite the larger trees associated with urban forests, overall stem density is lower, resulting in overall lower basal area and biomass. This may in turn be responsible for the lower canopy coverage in the forests with greater highway presence (Table 2), although the difference in coverage between the two extremes of urbanization was only 10%.

Compositional metrics were more variable in their influences than the structural metrics. Overall tree diversity was positively correlated with population density (Figure 5) and negatively correlated with mangrove and vegetation coverage (Table 2), which is consistent with terrestrial urban forests (Pickett et al., 2008). Mangrove diversity, however, showed the opposite effect and was highest in non-urban, unpopulated areas (Table 2) with greater open water and vegetation coverage (Figure 5). Little is known on the effect of urbanization on mangrove diversity, but other studies have pointed out the tendency of *L*. *racemosa* to form monoculture stands in post-disturbance shorelines (Benfield et al., 2005; Berger, Adams, Grimm, & Hildenbrandt, 2006). This agrees with our results, in which urban sites were dominated by *L*. *racemosa* (Figure 2), and the percentage of mangrove biomass as *L*. *racemosa* was most explained by the urban coverage within 1 km (Figure 5). Contrastingly, the contribution to mangrove biomass by *A*. *germinans* was negatively related to urbanization, suggesting that while *L*. *racemosa* is a synanthropic species, *A*. *germinans* is a misanthrop. Multiple regressions conflicted somewhat with simple regressions in these responses, and there was no significant simple regression to predict the contribution to biomass by *R*. *mangle*, suggesting more research is needed to produce more conclusive results.

Overall, the urban variables of urban cover, population density, and road and highway lengths explained 27% of the variation in forest structure and composition in the multiple regression models. The combination of all variables explained 67% of the variation of the data, on average, leaving 33% to be explained by other factors and randomness. Thus, urbanization is an important component of mangrove structure and function in Puerto Rico, as is to be expected from previous studies on both terrestrial and mangrove forests. Further, the importance of urban variables in some response metrics were only apparent in multiple regression, meaning all potential influences (e.g. water chemistry and hydrology) should be considered when evaluating urban mangrove structure and composition. Still, a previous study found only small influences of urbanization on hydrology and water chemistry in the same mangroves (B. L. Branoff, 2018), meaning the influence of urbanization was not fully detected in that study, or that the observed effect of urbanization seen in this study is due to other untested variables. Also, a sampling distance of fifty meters was the most commonly selected among the land cover variables, suggesting close range land cover is most important to mangrove forests.

As different kinds of mangrove forests provide different goods and services, so will the urban mangroves of Puerto Rico (Ewel, Twilley, & Ong, 1998). This is due not only to the variations in forest structure and composition, but also to the elevated number of people who serve to benefit from such services. Studies have suggested that higher stem density forests and forests of *Rhizophora* provide the greatest energy attenuation and thus shoreline protection (Alongi, 2008), which has been given the highest monetary value of mangrove services (Barbier et al., 2011). This would mean less protection from the urban mangroves of San Juan, which exhibited lower stem densities, basal areas, and biomass, and which were more likely to be dominated by *Laguncularia*. This is also true for the service of fisheries support, which is given the second highest monetary valuation and which is also greatly dependent upon *Rhizophora* and Avicennia, which were lacking in the most urban sites (Aburto-Oropeza et al., 2008; Rönnbäck, Troell, Kautsky, & Primavera, 1999). But a review on urban mangrove fish assemblages shows both benefits and drawbacks to urban land cover (B. Branoff, 2018). Other services include air and water filtration, microclimate regulation, noise reduction, and recreational services among others. These services depend upon many variables in terrestrial forests (Bolund & Hunhammar, 1999) but much less is known of these dependencies in urban mangrove forests and should be the focus of well-designed experiments.

Urbanization presents a particular challenge to tropical coastlines and especially Caribbean mangrove systems (Baird, 2009; McGranahan, Balk, & Anderson, 2007; Seto et al., 2012). Conversion of mangrove forests to developed lands has been observed to occur more quickly around some of the largest cities in the world (B. L. Branoff, 2017), thus urban mangrove conservation is an important priority for these systems. But understanding how the remaining forests function in their urban surroundings will also be an important component of management towards optimizing ecosystem services. This study showed that there are quantifiable patterns of forest change along an urban gradient and that these changes could mean a reduction in ecosystem services. Future work in these forests will assess the use of the mangroves by avifauna, anurans, and insects, as well as tree ecophysiological parameters and growth rates. But more studies are needed to support these results in the varying mangrove habitats of the world. These studies should be conducted along well defined and quantified urban gradients, allowing for the isolation of potential causal factors. Doing so will contribute to more informed management of mangroves in the Anthropocene.

